# Dengue and Chikungunya among Febrile Outpatients in Kinshasa, Democratic Republic of Congo: a cross-sectional study

**DOI:** 10.1101/486407

**Authors:** Sam Proesmans, Freddy Katshongo, John Milambu, Blaise Fungula, Hypolite Muhindo Mavoko, Steve Ahuka-Mundeke, Raquel Inocêncio da Luz, Marjan Van Esbroeck, Kevin K. Ariën, Lieselotte Cnops, Birgit De Smet, Pascal Lutumba, Jean-Pierre Van geertruyden, Veerle Vanlerberghe

## Abstract

**Background:** Pathogens causing acute fever, with the exception of malaria, remain largely unidentified in sub-Saharan Africa, given the local unavailability of diagnostic tests and the broad differential diagnosis.

**Methodology/Principal Findings:** We conducted a cross-sectional study including outpatient acute febrile syndromes in both children and adults, between November 2015 and June 2016 in Kinshasa, Democratic Republic of Congo. Serological and molecular diagnostic tests for arboviral infections were performed on blood, including PCR and NS1-RDT for acute, and ELISA and IFAT for past infections.

**Conclusions/Significance:** Among 342 patients, aged 2 to 68 years, 45.3% tested positive on malaria Rapid Diagnostic Test. However, 87.7% received antimalarial and 64.3% antibacterial treatment. Further investigation among 235 fever cases revealed 19 (8.1%) acute dengue and 2 (0.9%) acute chikungunya infections, with an important proportion of participants already exposed to flaviviridae (possibly dengue) and alphaviridae (possibly chikungunya) in the past, namely 30.2 % and 26.4% respectively. We found no evidence of exposure to Zika nor yellow fever virus.

**Author Summary:** Sam Proesmans is a medical doctor, currently doing a fellowship in internal medicine at the University of Antwerp, Belgium. He holds a Master of Public Health from Columbia University, New York City, USA. His primary research interest lies in infectious diseases, from a public health point of view and he believes that this study is proof that the differential diagnosis should include arboviral infection, given the hitherto unseen evidence of high arbivirus infection rates in the Democratic Republic of Congo, in that they are mostly misdiagnosed as malaria or bacterial infections.

## Introduction

Acute fever is one of the main reasons for healthcare seeking worldwide. In tropical settings, and especially sub-Saharan Africa, malaria is the first cause to be ruled out, which is done increasingly following the World Health Organization’s *testing before treating* policy of 2010 – through microscopic blood slide examination or rapid diagnostic test (RDT).

Following the introduction of this policy, together with the roll out of the highly efficacious artemisinin-combination as first-line malaria treatment and efficacious vector control, the overall malaria burden declined over the last decade. Accordingly, clinicians face a relatively higher number of malaria-negative patients for whom they do not have a clear diagnosis (1–3). In sub-Saharan Africa, where healthcare settings are often resource-limited, healthcare providers face the daunting challenge pinpointing the causing agent of these febrile illnesses in an adequate and timely fashion, with little to no diagnostic means other than a malaria-RDT. They mostly rely on history taking and physical examination to determine the focus and cause of infection, of which acute respiratory infection (ARI), gastroenteritis (GE) and urinary tract infection (UTI) are the three most prevalent syndromes reported. However, for over more than half of the patients presenting with acute fever, no focus can be found at all and clinicians label them as ‘undifferentiated’– knowing that their differential diagnosis is broad, ranging from viral, bacterial, parasitic to fungal infections. Although viral illnesses are often suspected, prescription of antimicrobials is rampant in this group (4) and their licentious usage in low-resource settings fuels the global burden of antimicrobial resistance (5–7). Thus more insight in the exact causes of this group of ‘undifferentiated fevers’ may help curb the usage of antimicrobials and improve the clinical care of patients in low-resource settings more broadly. Still, evidence on the causes of ‘undifferentiated acute fever syndromes’ is scarce and is coming from inpatient settings.

To our knowledge, only one study in sub-Saharan Africa (Sierra Leone) looked in a prospective way to etiologies of acute undifferentiated fever in all age groups at an outpatient, primary health care level and found a 5% dengue virus (DENV) infection rate and a 39% acute chikungunya virus (CHIKV) or related alphavirus infection rate (8). Nonetheless, limited outbreaks and sporadic clinical cases of DENV have been reported over the last 50 years in 22 African countries (9). Seroprevalence studies have demonstrated DENV IgG-antibodies, indicating past-infection, in 12.5% of study participants in Cameroon, 36% in Burkina Faso and 45% in Nigeria (9), although in other areas seropositivity remained zero (10). In Tanzania past infection rates are higher, reaching 50.6% in health-facility based studies and 11% in community-based studies (11). Despite the presence of all four DENV serotypes, severe disease epidemics are rarely reported in Africa (12). The DENV burden in Africa is, based on modeling, estimated at 16 million symptomatic clinical infections or 16% of the global total (13). In East-Africa, CHIKV outbreaks and circulation are described, such as in Kenya with a past-infection rate of 67% (14) and the reports of epidemics in 2004 in Kenya (15), in 2013 in Tanzania (16) and in 2018 in Mozambique (17).

In the Central African region, DENV, CHIKV, yellow fever (YFV) and Zika virus (ZIKV) – all leading causes of acute fever in other (sub)tropical regions – are circulating too. This is illustrated by CHIKV outbreaks in Kinshasa in 2000 (18) and 2012 (18bis) and Brazzaville in 2011 (19), and the 2013 DENV (20) and 2016 YFV outbreak in Angola (21), along with the first ZIKV case reported in Angola in 2017 (22). These outbreaks are only possible because the *Aedes* mosquito, the vector of the aforementioned arboviruses, thrives in this region. Furthermore, alphaviruses, flaviviruses and bunyaviruses were demonstrated to have infected *Aedes* and *Culex* mosquitoes in Kinshasa in 2014 (23).

In the Democratic Republic of Congo (DRC) the circulating pathogens causing uncomplicated acute undifferentiated fever, are unknown (24). However, outside the above documented epidemics, CHIKV and DENV probably circulate continuously. DRC travelers accounted for 22% of returned Africa travelers with a CHIKV infection in the outpatient clinic of the Institute of Tropical Medicine, Antwerp between 2007 and 2012 (25) and up to that point there was an increasing number of confirmed DENV infections in travelers coming from a large set of African countries, including DRC (26). In Eastern DRC, a few DENV cases have been found during an outbreak of West-Nile fever in 1998 (27) and between 2003 and 2012 when testing samples negative for YFV (28).

In this study in DRC, we aim to quantify the importance of arboviruses as a cause of acute febrile syndromes. Furthermore, we aim to describe the case presentation and the presence of arbovirus/malaria co-infections.

## Methods

### Setting

The study took place in Lisungi Health center in Pumbu, an area of about 14,000 inhabitants, belonging to the peri-urban health district Mont Ngafula 1, at the southern side of Kinshasa. The climate is tropical with a rainy season between October and May, and a dry season from June to September. The Lisungi health center is the only public health facility in the area, with an average of 250 consultations per week. Over the years, around 70% of the patients mention fever as the reason for consultation of whom half tested positive for malaria on RDT (personal communication with Dr. Blaise Fungula). The Lisungi health center is a higher-level clinic, has recently performed a *Good Clinical Laboratory Practice* compliant trial and has been involved in other febrile illness investigations, specifically on malaria (29).

### Study design and participants

The study was designed as a cross-sectional study with prospective patient inclusion. As the proportion of pathogens can change over time, especially for epidemic-prone diseases, we included patients proportionally from November 2015 to June 2016 and limited patient inclusion of undifferentiated fever to 6 per day (3 children and 3 adults). Only patients of at least 2 years old, presenting at the outpatient department with a history of acute fever (*i.e.* ≥ 2 days and ≤ 7 days) or having an axillary temperature of ≥ 37.5°C, were eligible. Patients with any history of an acute injury, trauma or poisoning, suspicion of meningitis/encephalitis, recent hospitalization or giving birth in the preceding two weeks, were excluded. Reported recent intake of antimicrobials was not an exclusion criterion, but was recorded accordingly. Given the non-specificity of the signs and symptoms of a possible arboviral infection, and in order to estimate the burden of co-infection, we included a subsample of patients with acute respiratory infection (ARI), gastroenteritis (GE) and urinary tract infection (UTI) categorized on clinical grounds or with confirmed malaria based on laboratory analysis. Only 1 child and 1 adult per day could be recruited from these latter three ‘differentiated fevers’ groups, in addition to the aforementioned maximum of 6 undifferentiated fever cases per day, in order to overcome selection bias. Our main interest was the distribution of viral pathogens among the undifferentiated fevers. To be able to detect the presence of a disease whose prevalence is 5% with a precision of 2.5% at a confidence level of 95%, 290 patients with ‘undifferentiated fever’ needed to be included. Increased with 10% for incomplete data or loss of biological samples, we came to a sample size of 320.

### Data collection

#### Case Report Form

For every participant, a pre-tested case report form (CRF) was filled out, based on a standardized clinical history and physical examination by one of the four physicians/clinical officers on ground, following standard clinical practice. Prior to the start of study, the clinicians received a training on the inclusion algorithm and CRF. Patient management was done according to local guidelines. For every patient the following information was recorded: demographic data (age, sex, residence), past medical history (yellow fever vaccination, intake of antimicrobials), signs and symptoms, duration of fever, medical examination, diagnosis and treatment given.

#### Laboratory analyses

In the on-site laboratory of the Lisungi health center, capillary blood was taken to microscopically count the number of leukocytes and a thin smear stained with Giemsa was done for differentiation. Hematocrit and hemoglobin were determined using a portable spectrophotometer (Hemocontrol, EKF Diagnostics, Barleben, Germany). Furthermore, malaria and dengue were diagnosed by the HRP2/pLDH RDT SD Bioline® Malaria Ag Pf/Pan 05FK60 and RDT SD Bioline® Dengue Duo 11FK46 respectively. The latter is an immunochromatographic one-step assay designed to detect both DENV NS1 antigen (present in first week of infection) and IgM/IgG antibodies (detectable from the fifth day of fever onwards) to DENV. HIV was not routinely tested for, given the overall low prevalence in the study area and the unavailability of HIV antiretroviral treatment according to national ethics guidelines in this setting.

All undifferentiated fevers, together with about half of both the malaria positives and the clinically apparent differentiated fevers - identified through random selection, were tested for arbovirus exposure. For this purpose serum and EDTA whole blood were stored in liquid nitrogen (−196°C) on-site and transported afterwards to the Institut National de Recherche Biomédicale (INRB) in Kinshasa. After six and nine months, the samples were airlifted to the Institute of Tropical Medicine in Antwerp, Belgium (ITM) for additional arbovirus specific tests. Molecular tests were done for DENV, CHIKV and YFV on all available samples. For ZIKV, as there was no evidence of its circulation in the area, testing was done on 50 randomly selected undifferentiated fevers and 30 ‘clinically apparent differentiated fevers’ (through the random numbers tool in Microsoft Excel). The real-time reverse transcriptase polymerase-chain reaction (RT-PCR) for DENV, CHIKV, ZIKV and YFV were done with in house PCR protocols currently in use in the reference laboratory of ITM. The IgM and IgG antibodies for DENV and CHIKV were detected using enzyme-linked immunosorbent assays (ELISA) (Dengue Virus IgM Capture DxSelectTM and Dengue Virus IgG DxSelectTM from Focus Diagnostics, Cypress, CA, USA and CHIKV ELISA IgM and CHIKV ELISA IgG test from Euroimmun, Lübeck, Germany). An indirect immunofluorescence antibody technique (anti-CHIKV indirect immunofluorescent test (IIFT) from Euroimmun, Lübeck, Germany) for both CHIKV IgM and IgG was done on the ELISA positive samples and a random selection of 20 negative samples. In case of a non-congruent result between ELISA and IIFT, a malaria PCR was done to evaluate if cross-reaction between malaria and CHIKV could explain this finding (30). A selection of IgG DENV ELISA positive samples were tested with DENV and YFV plaque reduction neutralization tests (PRNT) to differentiate between past exposure to DENV or vaccination against YFV.

#### Case definitions

Acute DENV infection was defined as a positive DENV-PCR and/or a positive DENV-IgM-ELISA.

Past DENV infection was defined as a positive DENV-IgG-ELISA in the absence of DENV-NS1, RNA and IgM antibodies.

Acute CHIKV infection was defined as a positive CHIKV-PCR and/or positive CHIKV-IgM by IIFT.

Past CHIKV infection was defined as a positive CHIKV-IgG by IIFT in the absence of CHIKV RNA and IgM antibodies.

Acute YFV infection was defined by a positive YFV-RT-PCR. Acute ZIKV infection was defined by a positive ZIKV-RT-PCR.

#### Statistical analysis

Data were entered into an Excel database (Microsoft Corp, Va., USA). Statistical analyses were performed with SPSS Statistics version 23 (IBM, NY, USA). The Pearson Chi-square test was used to determine the association between categorical variables. An alpha level of 0.05 was used for all tests for statistical significance.

### Ethics statement

This study was approved by the ethical review boards of the School of Public Health of the University of Kinshasa (DRC), the Institute of Tropical Medicine of Antwerp (Belgium) and the University Hospital of Antwerp (Belgium). The study was registered in a public repository (https://www.clinicaltrials.gov/ct2/show/NCT02656862). Written informed consent was obtained from every adult or – in case of minors – from their caretaker. This study was conducted in full compliance with the principles of the latest amended *Declaration of Helsinki* and of the *International Conference Harmonization* (ICH) guidelines, plus adhering to local laws and regulations. This study was registered in the clinical trial register under the number: NCT02656862.

## Results

Over the period November 2015 to June 2016, 342 patients were included from whom clinical data and on the spot malaria-RDT results were recorded. Clinical diagnoses were as follows: 70.2% undifferentiated fever, 6.7% UTI, 6.1% GE and 17.0% ARI - all three further labeled as ‘clinically apparent differentiated fevers’. The study population (Table 1) consisted of 183 (53.5%) female participants and 180 (53.1%) were under the age of 18. Fifty (14.6%) patients were in need of hospitalization. The reported YFV vaccination rate was as low as 1%. The vast majority (77.8%) of the patients presented in the four first days after the onset of fever, with relatively nonspecific signs and symptoms (Table 2). Malaria RDT was positive in 155 (45.3%) of the participants. However, 300 (87.7%) received an antimalarial treatment. Antibacterials were frequently prescribed: 64.3% of participants received at least one and 11.7% received even more than one antibacterial drug.

**Table 1.** Baseline data of acute fever patients and their arbovirus exposure at Lisungi Health, DRCongo, 11/2015-06/2016 (n=342, n tested=235)

| Characteristics | Total<br>number<br>N=342 (%) | Recent<br>dengue<br>infection<br>(n=19)<br>N (%) | Recent<br>chikungun<br>ya<br>infection<br>(n= 2 )<br>N (%) | Past<br>arbovirus<br>infection<br>(n=71)<br>N (%) | Past<br>alphavirus<br>infection<br>(n=62)<br>N (%) |
| --- | --- | --- | --- | --- | --- |
| Age groups<br>(years): |  |  |  |  |  |
| < 5 | 56 (16.5) | 3 (15.8) | 0 | 7 (9.9) | 2 (3.3) |
| 5 - 17 | 124 (36.6) | 6 (31.6) | 0 | 10 (14.1) | 16 (26.7) |
| 18 - 44 | 115 (33.9) | 8 (42.1) | 2 (100) | 35 (49.3) | 27 (45.0) |
| 45 - 64 | 39 (11.5) | 2 (10.5) | 0 | 15 (21.1) | 13 (21.7) |
| ≥ 65 | 5 (1.5) | 0 | 0 | 4 (5.6) | 2 (3.3) |
| unknown | 3 | 0 | 0 | 0 | 2 |
| Gender, Female | 183 (53.5) | 11 (57.9) | 2 (100) | 36 (50.7) | 31 (50.0) |
| Consulting during<br>dry season | 52 (15.2) | 10 (52.6) | 0 (0) | 10 (14.1) | 12 (19.4) |
| Commune of<br>residence: |  |  |  |  |  |
| Lemba | 2 (0.6) | 0 | 0 | 0 | 0 |
| Makala | 1 (0.3) | 0 | 0 | 0 | 0 |
| Matete | 1 (0.3) | 0 | 0 | 0 | 0 |
| Mont-Ngafula | 250 (73.1) | 16 (84.2) | 2 (100) | 49 (69.0) | 50 (80.6) |
| Ngaliema | 32 (9.4) | 2(10.5) | 0 | 5 (7.0) | 6 (9.7) |
| Selembao | 56 (16.4) | 1 (5.3) | 0 | 17 (23.9) | 6 (9.7) |
| Recent travel | 10 (2.9) | 1 (5.3) | 0 (0) | 4 (5.6) | 1 (1.6) |
| Reported yellow<br>fever vaccination | 4 (1.2) | 0 (0) | 0 (0) | 1 (1.4) | 0 (0) |

**Table 2.**
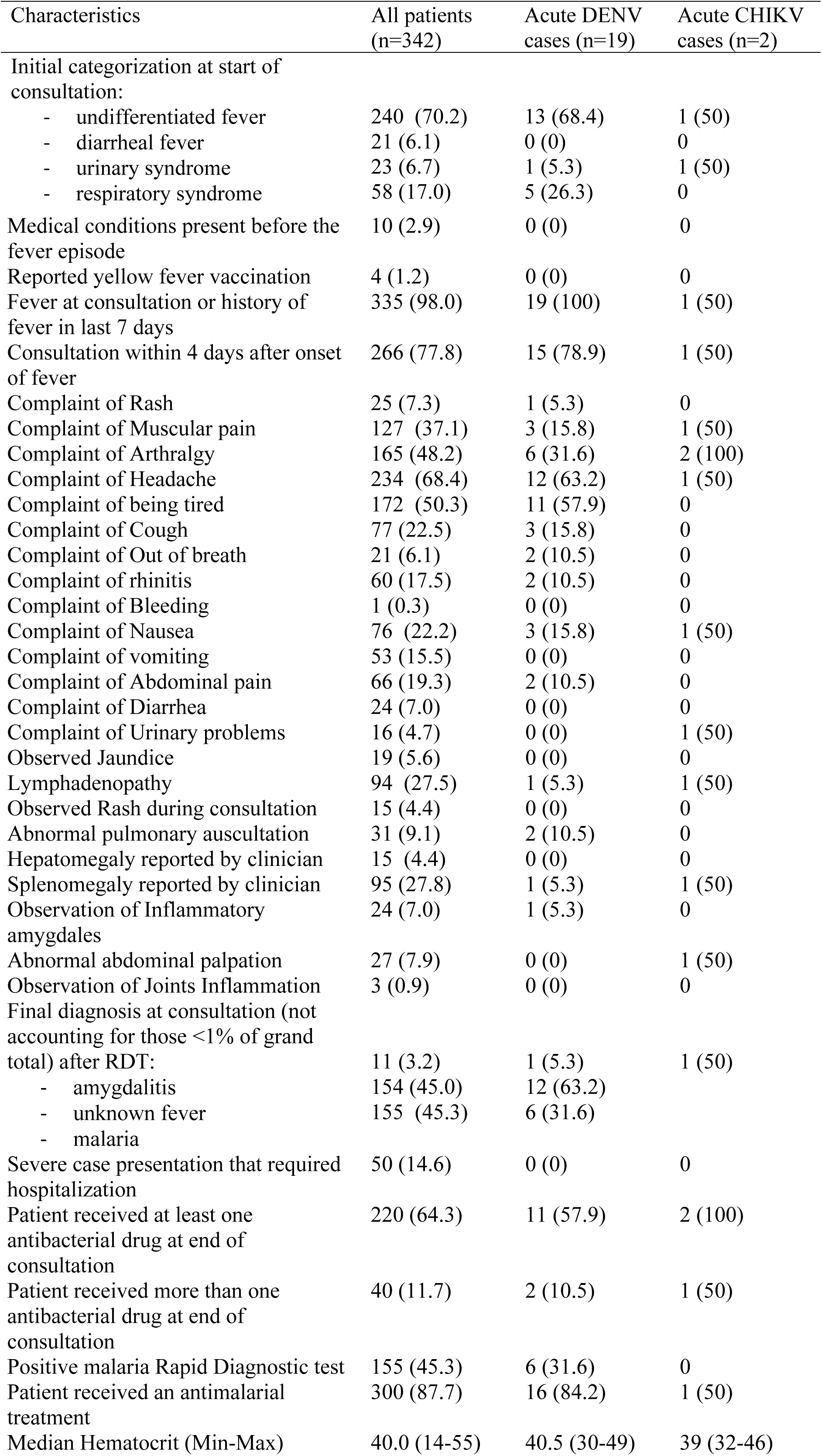
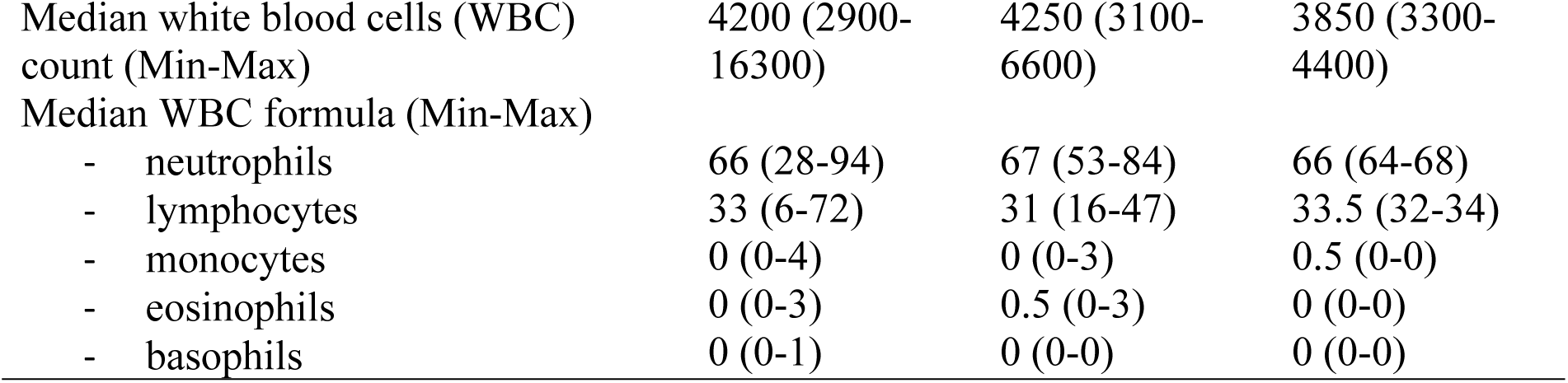
Clinical characteristics of acute fever patients stratified by DENV and CHIKV infection at Lisungi Health, DRCongo, 11/2015-06/2016.

In the subsample of 235 further tested for arboviruses, 19 (8.1%) fulfilled the criteria of an acute DENV infection, of which 14 were confirmed by PCR. Both DENV1 and DENV2 were detected (Table 3). In contrast, only two acute CHIKV infections were diagnosed, both by the presence of IgM antibodies. The majority of CHIKV IgM ELISA positive samples was not confirmed by IIFT. On these non-congruent samples, PCR to detect Plasmodium was performed and revealed an actual malaria infection in 18 of the 22 positive IgM ELISA - negative IgM IIFT. In the dry season, there was an increased risk of 7.2 (95% CI 2.6 - 20.0) of presenting with acute DENV in comparison to the rainy season (Supporting Information file 1). Of the acute DENV cases, 31.6% had a positive malaria RDT versus 37.0% among the other causes of fever (p=0.64), stressing the importance of co-infection. Furthermore, of the malaria negative cases, 8.7% (13/149) tested positive for acute DENV. None of the acute CHIKV cases tested positive for malaria or vice versa. Neither acute ZIKV, nor YFV infections were detected.

**Table 3.**
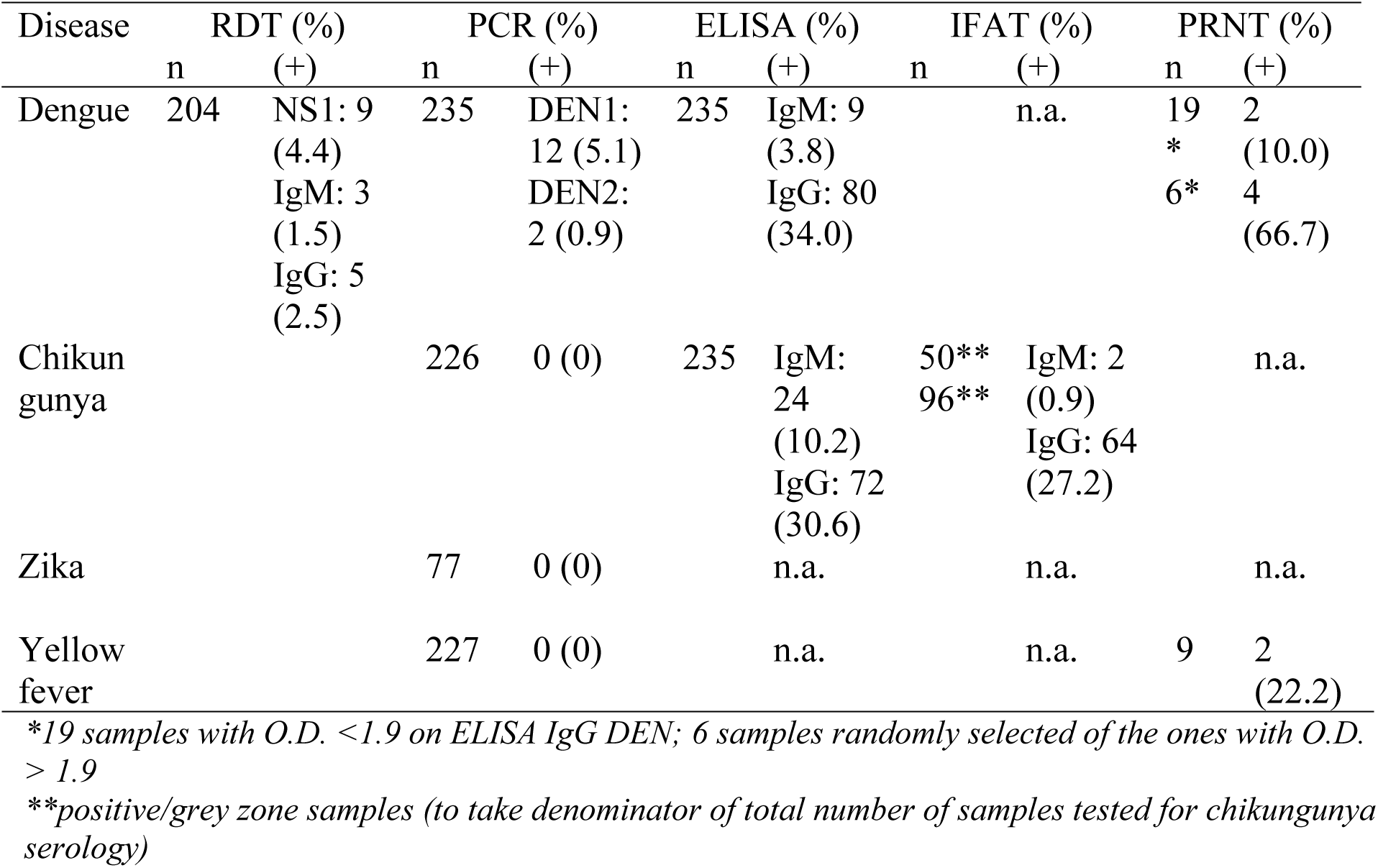
Results diagnostic laboratory tests for arbovirus infections at Lisungi Health, DRCongo, 11/2015-06/2016.

With regard to the clinical presentation of both DENV and CHIKV infections, we found no specific signs or symptoms to be statistically significantly – let alone clinically relevant – associated with acute DENV or CHIKV versus the other febrile patients (S1 Table). None of the acute DENV or CHIKV cases were severe, none were hospitalized and there was no apparent leucopenia or hemoconcentration (as often seen in severe DENV cases).

We found 71 (30.2%) patients with evidence for past DENV infection (ELISA-IgG positive), of which 60 (75%) contained relatively high levels of anti-flavivirus antibody (ratio ≥ 1.9), the latter not depending on age (data not shown). The PRNT on the subsample was only positive in 5.6% and 66.7% for DENV, and 25% and 20% for YFV, on samples with IgG ratio below 1.9 and above 1.9 respectively. For these past infections, all 4 serotypes were detected: DENV1, DENV2, DENV3 and DENV4 in 4, 2, 1 and 1 patient respectively (of the 6 positively tested DENV PRNT) – two patients were reactive to all 4 serotypes, the other 4 only to 1 serotype. Past exposure to CHIKV was confirmed with IFAT IgG in 26.4% of the study participants. When taking past DENV and CHIKV together, 56.6% of the study participants were exposed to at least one and 11.9% to both arboviruses. The prevalence of past DENV and CHIKV infections increased with age, raising from 18.9% and 5.4% under 5 years of age, to 80% and 40% over 65 years of age, respectively (Fig 1). The association of age is statistically significant for the past infections with CHIKV (p=0.01) and DENV (p<0.01) (S2 Table). Having been exposed to DENV was also statistically significant associated with recent travel (p=0.02). The congruence between the RDT and PCR/ELISA results for DENV was variable: in comparison to PCR the sensitivity of NS1 was 90%, in comparison to IgM ELISA the IgM RDT had a sensitivity of 30% and in comparison to IgG ELISA the IgG RDT had a sensitivity of 7.6%. The specificities were all above 99.3%.

**Figure 1.**
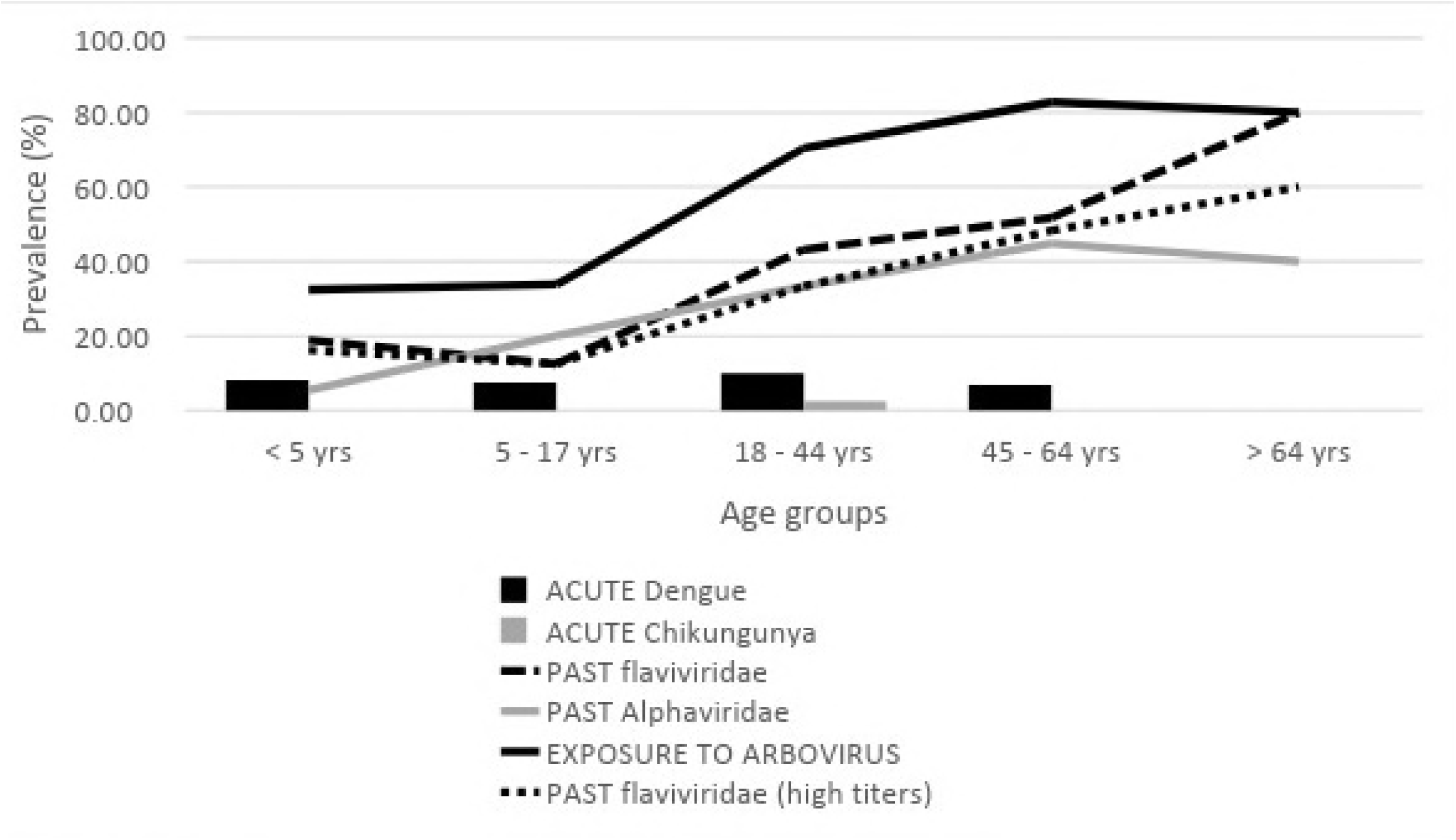
Prevalence of Flavi and Alpha viridae infections and their related seroprevalence in febrile patients stratified by age at Lisungi Health, DRCongo, 11/2015-06/2016 (n= 235)

## Discussion

Although no epidemics were reported recently, our study shows ongoing transmission of arboviruses in Kinshasa, DRC. Acute and past DENV infection was demonstrated in 8.1% and 30.2% of the participants, respectively. This is on the higher end of the spectrum compared to other studies in Africa reporting an overall flavivirus seroprevalence ranging between 0 and 35% with a mean of 18.1% (9,24). The IgG-seroprevalence increased with age suggesting a continuous exposure to the flaviviridae over time. The findings on CHIKV, 0.9% acute and 26.4% past infection rates respectively, were in line with the estimated seroprevalence of 34.4% in Congo Brazzaville before the outbreak of 2011 (19) and were on par with other African sites reporting an overall alphavirus seroprevalence oscillating between 0 and 72% (24). Remarkably, CHIKV IgG was also detected in small children, born after the 2011 epidemic, pointing towards an endemic circulation of the virus.

Importantly, neither DENV nor CHIKV was clinically suspected or was considered in the clinical differential diagnosis and 64.3% of patients were treated with at least one antibacterial drug, of whom almost one in six (11.7%) received dual or triple antimicrobial therapy. A possible explanation of the apparent absence of clinical and/or severe DENV cases in our study, but also in other African settings, is that African heritage genetically protects against severe DENV, more specifically the lower OSBPL10 expression profile in Africans is protective against viral hemorrhagic fever and dengue shock syndrome (31,32).

We used DENV ELISA tests on all samples. Although it is widely known that there is cross-reactivity among flaviviruses, ELISA is still the most affordable – hence most commonly used – test (24). Since YFV vaccination is an expected cause of cross-reactions, we performed PRNT for DENV and YFV in a subset of samples, and found that the congruence between DENV ELISA and PRNT was lower than in American and Asian settings (33–37). A similar observation was made in another study in DRC with a certain level of discordance between ELISA and PRNT (38), and in a study in Ethiopia where cross-reaction and low confirmation by PRNT was also shown (39). In our study, the majority of the samples negative with DENV PRNT was also negative with PRNT for YFV, which indicates that the high flavivirus IgG positivity was not due to YFV vaccination. A plausible explanation can be that in DRC other flaviviruses circulate, which cause the aforementioned cross-reaction in ELISA (40). Similarly, the results of CHIKV serology could be the result of cross-reaction with other togaviruses, but this is very unlikely in DRC as Mayaro is only present in South America, and there is no apparent circulation of o’nyong nyong, Semliki Forest and Sindbis virus. YFV was detected in mosquitoes collected in Kinshasa a year before our study (23) and was actively circulating in the region at the time of study, transgressing the border with Angola (21), but we did not detect any acute case. The report of YFV vaccination was very low (below 1%), but as YFV vaccination is included in the childhood vaccination program over the last decade, this may indicate that the population is not aware of which vaccines their children get. However, the increasing prevalence of flavivirus IgG antibodies with age is congruent with a history of increasing exposure to the pathogens over lifetime and thus was in this study not explained by YFV vaccination. ZIKV was not suspected to be circulating in the area and indeed, no positive case was found in the subsample tested.

In a recent study conducted in the same area in Kinshasa, it was reported that 62% of patients with acute fever had neither malaria nor unspecified bacterial infection (41). For the first time we were able to demonstrate the fact that arboviruses, more specifically DENV and CHIKV, circulate in high numbers in the capital of DRC. The highest number of cases were reported in the dry season, but cases were also confirmed in the other months, indicating that transmission is not only happening in epidemics, but that there is a year-round circulation.

This contradicts the existing paradigm that arbovirus infections are outbreak-driven in Africa and are not endemic, as is the case in Asia and Latin America. Our study adds to mounting - but still scarce – evidence that arboviruses are endemic in large parts of Africa (17).

Although the sampling design of this study was adequate to evaluate the proportion of arboviruses causing acute febrile syndromes, sample size was small and patients were only recruited from a single health center. However, the Lisungi health center is well visited by the surrounding population and all ages were represented in our study population. Moreover, the median age of our study population is 17 years, which approximates the median age of 18.6 years in the DRC (UNDESA 2017 and CIA World Factbook 2017).

The discrepant results between CHIKV ELISA and IIFT were at least partly explained by the cross-reaction with actual malaria infection, a phenomenon which has been described before for ZIKV (42). Furthermore, we were able to document the common practice of overprescription of antimalarial drugs in malaria RDT-negative patients, as is apparently the case nationwide in DRC as recently shown by Ntamabyaliro et al (43). Indeed, while not even half of the patients tested positive for malaria, almost 90% received antimalarial treatment, in addition to 16% of patients treated with over-the-counter antimalarials prior to presentation at the clinic, which brings the total close to 100%. It could be questioned whether the rigorous implementation and usage of RDTs has any added benefit. A recent meta-analysis evaluating data from over half a million children and adults, showed that the introduction of a malaria RDT simply shifted the antimicrobial overuse from one antimicrobial class to the other, mainly from antimalarial to antibacterial and anthelmintic drugs (44). Consequently, the increasing prescription rate of antimicrobials – including antibacterial, anthelmintic and antimalarial drugs, is extremely worrisome in terms of the growing global problem of antimicrobial resistance, including against *Plasmodium falciparum*.

We did not investigate bacterial causes of fever through culture of blood or other bodily fluids, a test typically done in hospital settings, hereby possibly underestimating the burden of concomitant bacterial (super)infection. We therefore encourage further research elucidating the broad range of pathogens causing febrile illness and the distribution of the vectors involved in arboviral infections in urban and rural sub-Saharan African settings.

Based on our findings, we recommend to include arboviral infections, namely DENV and CHIKV, in the differential diagnoses of acute fever presentation in sub-Saharan Africa.

## Acknowledgments

We would like to express our sincere gratitude to all the people, caregivers and beyond, who helped facilitate and conduct this study in Lisungi, Kinshasa and to Ms. Nikki Foqué for her excellent lab support at the Institute of Tropical Medicine, Antwerp.

## Supporting Information Legends

Supporting Information Table 1: factors associated with acute dengue infection

Supporting Information Table 2: Factors associated with past infection arbo- and alphaviruses S1 Checklist: STROBE Checklist

